# Back to the horns: a reconstruction of the ancestral horn state through distinct types of recombination events

**DOI:** 10.1101/2024.11.18.624115

**Authors:** Maulik Upadhyay, Alexander Graf, Neza Pogorevc, Doris Seichter, Ingolf Russ, Stefan Krebs, Saskia Meier, Ivica Medugorac

**Author notes:** Corresponding author: IM.

## Abstract

**Background:** Breeding of the genetically polled animals is the desirable approach in modern cattle husbandry. At least four different genetic variants associated with polledness in cattle have been identified, suggesting genetic heterogeneity. These dominant variants have been identified on chromosome 1 between the regions of approx. 2.42 to 2.73 Mb (reference: ARS-UCD1.2), also called the *POLLED* locus. Among these variants, Friesian (*P_F_*, 80 kbp duplication) and Celtic (*P_C_*, 212 bp complex InDel) are the most observed in the majority of breeds in the production systems globally, such as in Holstein-Friesian (HF) and Fleckvieh (FV). While a putative causal association of *P_C_* with polledness is proven, the presence of large duplication in *P_F_* makes it difficult to prove the causality.

**Results:** In this study, we conduct whole-genome sequencing (WGS) analysis of two trios exhibiting unexpected inheritance patterns related to *P_C_* and *P_F_* variants. In both instances, horned offspring were produced from mating pairs where one parent was homozygous for the polled variant, and the other was homozygous for the ancestral horned variant. By analyzing the WGS data generated using long-read sequence technology, we show that *de-novo* generation of the ancestral horned variant in both the offspring was the result of distinct recombination events. Specifically, in case of the HF trio, it was the result of non-allelic homologous recombination in the gametes of the sire (*P_F_/P_F_)*, while in case of the FV trio, it was the result of allelic homologous recombination in the gametes of the dam (*P_C_/P_F_*). The findings from the HF trio support the hypothesis that the 80-kbp duplication is the genetic variant responsible for the polled phenotype of Friesian origin.

**Conclusion:** Here we show that different genomic arrangements in *POLLED* locus can lead to the emergence of *de-novo* ancestral horn phenotypes. These kinds of arrangements can make a reliable gene test less reliable and the derivation of the phenotype difficult to predict. Therefore, it is important for the large *POLLED*-locus that any deviation from the expected result is critically analyzed. Possibly some of these cases can further narrow down the sequence motif that is essential for polledness in cattle.

## Background

Based on the size, shape, and characteristics of horns, the head phenotype in cattle can be categorized into these three classes: (i) true horns in bovids contain a pneumatized bony inner core that is attached to the frontal bone and have an outer keratin sheath [1]; (ii) scurs are rudimentary horn-like appendages that are loosely attached to the skin rather than the frontal bone with not pneumatized bony inner core. (iii) complete absence of horns-like appendages, a condition called polledness. Like their wild ancestors, the aurochs, and all members of the Bovidae family, cattle are naturally horned. Therefore, the presence of true horns is an ancestral characteristic of Bovidae, and scurs and polledness are derived traits. Horns have been associated with a diverse range of functions such as in exerting territorial dominance, competing for mating and other resources, protecting against predators, and maintaining thermoregulation [2–4]. The process of domestication and subsequent breed formation has largely rendered horns redundant. In fact, in modern industrial settings, horned animals are undesirable as they pose injury risks to themselves, other cattle as well as their handlers. Further, it is commonly assumed that because of its special requirements, horned cattle are associated with high barn investment and labor costs [5]. Therefore, polled conditions in cattle have become highly desirable. To this end, a variety of chemical and surgical methods are used for disbudding or dehorning. Across the cattle sector, these practices are commonplace. According to a survey published in 2015, in Europe, more than 80% of dairy, 46% of beef, and 67% of suckler calves are disbudded or dehorned [6]. All these procedures are labor intensive and cause pain and distress to animals; further, they also leave the wound sites exposed to infection, adding to the distress [5, 7]. Therefore, the animal welfare guidelines recommend adapting husbandry systems to the ancestral characteristics of cattle or breeding genetically hornless animals rather than dehorning them.

Early and accurate detection of animals having genetic predisposition for polled phenotype is key to selective breeding for polled animals [8]. Polled phenotype is a Mendelian trait governed by multiple variants (genetic heterogeneity) that are located on BTA1 (bovine autosome 1) between the region of approx. 2.42 to 2.73 Mb (reference: ARS-UCD1.2), also referred to as *POLLED* locus throughout the manuscript. Phenotypic penetration suggests that all these variants are dominant and cause polledness. Mainly, these four polled variants have been identified so far: (i) Celtic (*P_C_*): This complex insertion-deletion variant, “g.[2429326_2429335del;2429109_2429320dupins]”, was first identified in several European breeds; since the geographical distribution of these breeds overlaps mainly with the area in which Celtic culture was practiced, it was named as “Celtic” [9]. (ii). Friesian (*P_F_*): this tandem duplication variant, “g.2629116_2709243dup (80,128 bp)”, was first identified in Holstein-Friesian (HF) cattle [9, 10]. The duplicated segment has the same orientation as the reference sequence. It differs from the reference sequence by a SNP (single Nucleotie polymorphism) (*P_T->A_)* and 2 bp deletion (*P_F2D_,* TG) on 1’st bp and 38th bp positions respectively on the duplicated sequence. (iii). Mongolian (*P_M_*): this complex insertion-deletion variant, “[2695261_2695267delinsTCTGAA;2695889_2696047dupins]”, was first identified in Mongolian Turano cattle and Mongolian yaks [11]. The *P_M_* variant is located within the 80,128 bp segment duplicated by *P_F_* duplication. (iv). Gaurani (*P_G_*): This duplication variant, “g.2614828_2724315dup”, was first identified in Nellore cattle from Brazil [12] and completely overlaps with *P_F_* and *P_M_*. Interestingly, none of these four variants are directly involved in the gene coding. While several hypotheses have been proposed [13, 14], the exact pathways through which these complex variants disrupt the migration or proliferation or differentiation of cells in the horn bud region is not known so far [15].

Among these four variants, *P_C_* and *P_F_* are the most observed polled variants in the majority of breeds in the production systems globally. Of these variants, a putative causal association of *P_C_* with polledness is proven by introgression of *P_C_* allele from genome-edited cell lines resulting in the birth of polled calves [16]. For the *P_F_* variant, the presence of large duplication (∼80 kb) makes it difficult to prove the causality by genome editing or to pinpoint the exact genomic region responsible for the polledness condition. The relatively small *P_M_* structural variant (SV), that is embedded in *P_F_*, however, suggests that the more distal part of the 80 kb region could be essential to the polledness. The question still remains whether any SV along the entire *POLLED* locus (i.e. from the start of the *P_C_* duplication (2,429,326) to the end of the *P_G_* segment (2,724,315) is sufficient to disrupt the process of horn bud formation.

In this study, we conducted whole-genome sequencing (WGS) analysis of two trios exhibiting unexpected inheritance patterns related to *P_C_* and *P_F_* variants. In both instances, horned offspring were produced from mating pairs where one parent was homozygous for the polled variant, and the other was homozygous for the ancestral horned variant. The polled gene test targeting 2-bp deletion (*P_F2D_*) and 212 bp insertion (entire *P_C_* variant) was used to confirm the presence of *P_F_* and *P_C_* variants, respectively. Specifically, in the HF trio, the sire was homozygous *P_F_*/*P_F_*, while in the fleckvieh (FV) trio, the dam carried *P_C_*/*P_F_* genotype. Therefore, resulting offspring were expected to exhibit polledness by inheriting either *P_C_* or *P_F_* polled variant. To our surprise, here investigated offspring from each trio, showed the regular horn condition and *p*/*p* genotype instead. After confirming parentage with BovineSNP50 BeadChip, we hypothesized that recombination events in the *POLLED* locus of the paternal germ cells in HF trio and maternal germ cells in FV trio may have resulted in a novel genomic rearrangement, giving rise to the variant similar to the wild type. These trios provided us an opportunity to shed light on the genomic composition of the *P_C_* and *P_F_*variants.

## Methods

### Sample information

In this study, we investigated two trios, one from the HF breed and one from the FV breed. In the HF trio, the sire was confirmed to be homozygous for the *P_F_* variant, while the dam expressed a horned phenotype and according to genetic test *p*/*p* genotype. Notably, the sire has over 13,000 registered offspring, of which only one examined here was declared to exhibit the horned phenotype. In the FV trio, the dam carried one copy of both the *P_C_* and *P_F_* variants, while the sire was tested as horned with *p/p* genotype. The presence of the *P_F_* variant in the FV population was expected, as this breed is, since 1960s, introgressed by HF and thus carries a minor but significant genetic ancestry from the HF breed. In both trios, offspring were genotyped as *p/p* and exhibited regular horns fused to frontal bone instead of expected polled phenotypes and polled genotypes.

### DNA extraction and whole genome long-read sequencing

The DNA was extracted from the frozen blood or semen (sires) samples immediately before the sequencing to get an increased fraction of long DNA fragments in the sequencing. The DNA extraction was carried out from 100 µl blood using the QiaAmp Blood Mini Kit kit protocol (Qiagen, Hilden, Germany). Briefly, the resuspended blood was lysed and digested using proteinase K and RNase A, followed by binding to a column, washing, and elution through centrifugation.

To carry out long-read sequencing using the Oxford PromethION sequencer, one µg of unsheared DNA was end-repaired and A-tailed with NEBnext UltraII End-repair and A-tailing module (New England Biolabs, Ipswich USA). According to the manufacturer’s protocols, this mixture was later purified with AmpureXP magnetic beads (Beckman Coulter, Brea USA), ligated to 1D sequencing adapters (LSK114 kit, Oxford Nanopore Technologies (ONT), Oxford, UK), and again purified with AmpureXP beads. In the last Ampure cleanup step, instead of washing with 70% ethanol, a washing buffer supplied by Oxford Nanopore Technologies (LSK114 kit) was used. Approximately 200 ng of adapted DNA library was loaded to a primed PromethION R10.4.1M flow cell and sequenced for 72 h on a PromethION P24 sequencer.

### SNP genotyping, haplotyping and parentage and Polled test

Both trios were genotyped with the Illumina BovineSNP50 BeadChip (Illumina San Diego, USA) according to the manufacturer’s protocol. All 54,157 markers were used to test the parentage in both trios. Subsequently, we confirmed the expected pedigree in both the trios and for FV trio, even traced the pedigree to much deeper level than the first generation pedigree because our genotypic database included four FV grandparents. To investigate the recombination events, the inference of haplotypes and imputation of missing genotypes were carried out using hidden Markov models as implemented in BEAGLE (v 5.0) [17, 18]. In order to improve the accuracy of the phasing, genotype and pedigree information of additional trios and pairs of both breeds genotyped with BovineSNP50K BeadChip, otherwise not included in the analysis of this paper, were also included in this step of phasing.

Polled test was carried out according to the procedure and primers described in Medugorac *et al*.[9]. Briefly, the primer binding sites flanking the *P_C_* variant InDel events (5’- TCAAGAAGGCGGCACTATCT-3’ and 5’-TGATAAACTGACCCTCTGCCTATA-3’) were PCR amplified (94 °C 30 sec, 58 °C 60 sec, 72 °C 60 sec for 31 cycles). Next, PCR products were size-separated and visualized using 2% ethidium-bromide stained agarose gel electrophoresis. Genotyping of the *P_F_* 80 kb duplication event was carried out using two primers flanking the variable site *P_F2D_* (5’-GAAGTCGGTGGTCTGAAAGG-3’ and 5’- TGTTCTGTGTGGGTTTGAGG-3’). Next, PCR amplification (32 cycles 94 °C 30 sec, 59 °C 60 sec, 72 °C 60 sec) generated a RefSeq related product (*p_ref_*); this product was obtained in all animals, be it horned or polled. Additionally, a second product related to *P_F_* variant, containing the two-base pair (TG) deletion was only observed in the animals that carried one or two copies of the *P_F_* variant (*P_FDUP_*), see (Figure 1). To discriminate between these two products related to *P_F_* variant, ABI Prism 3130XL DNA sequencer was used. Animals carrying *P_F_/P_F_* were distinguished from *P_F_/p* using quantitative evaluation of the obtained signals. The animals carrying *P_F_/P_F_*yielded signals of similar peak heights as the *P_F_* and *p_ref_* products, while heterozygous products (*P_F_/p*) resulted in signal intensities of approximately double height for the p_ref_ product when compared to the product specific to *P_F_*.

**Figure 1.**
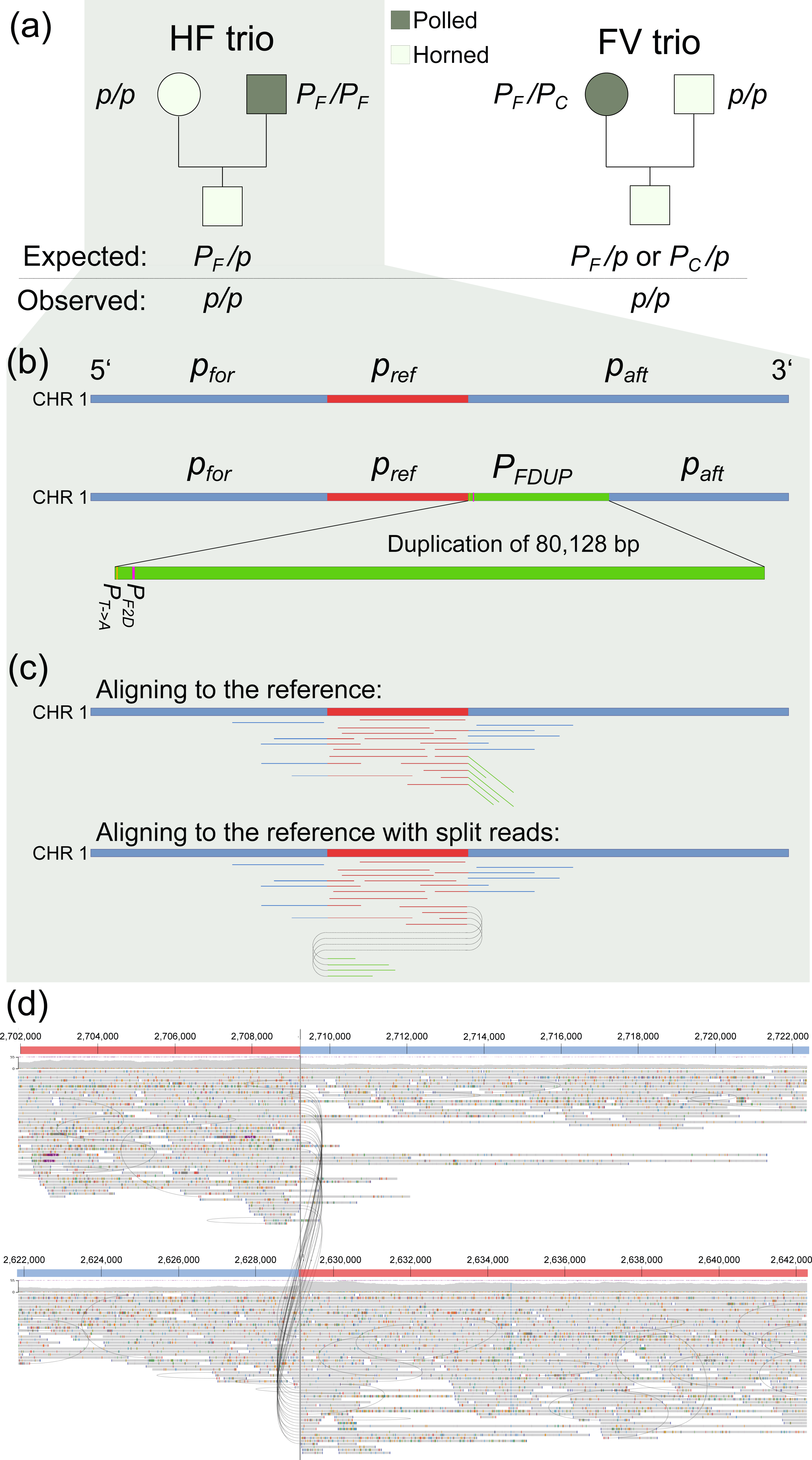
Overview of the inheritance patterns in the trios and genomic composition of the *P_F_* variant. (a) a schematic representation of the trios, including their genotypes, (b) a schematic representation of the wild type locus and friesian polled variant (*P_F_*); note that *P_F_* variant consists of a reference segment *p_ref_* and its duplication *P_FDUP_* segment. Two variants, namely, *P_T->A_*, *P_F2D_*, depicted on the 5’ side of *P_FDUP_* segment refer to a SNP and a 2 bp InDel already reported to be present on *P_FDUP_*, (c) a schematic representation of the patterns in which the reads sequenced from a sample containing *P_F_* variant would align to the reference, (d) actual screenshot of the split reads sequenced from the HF sire which is homozygous for *P_F_*variant.

### Bioinformatics pipeline to analyze WGS data

The samples were sequenced on a PromethION sequencer from Oxford Nanopore and were basecalled and demultiplexed during sequencing using Dorado (Dorado basecall server: v.7.3.11+0112dde09). After basecalling, basic quality checks of the sequences were carried out using NanoPlot (v 1.41.6) [19]. Subsequently, the sequencing data were aligned to the cattle reference genome sequence (ARS-UCD1.2) using minimap2 (v 2.28) with secondary alignments suppressed [20, 21]. Small genomic variants (SNPs and short InDels) were identified using Clair3 (v 1.0.10) with the appropriate trained model for the basecaller. Structural variations (SVs) were detected using Sniffles2 (v 2.4), cuteSV (v 2.1.1), Dysgu (v 1.6.7) and SVIM (v 2.0.0)[22–26]. Moreover, SVs were also detected using a de novo genome assembly-based approach. For this purpose, Flye (v 2.9.5) [27] was used for de novo assembly of the trios, and SVIM-asm (v 1.3.0) [28] was used for the detection of SVs. Finally, all genomic variants were phased using LongPhase (v 1.7.3) [29].

To investigate the split-read alignments in the POLLED locus region and to identify recombination breakpoints in the offspring’s genome using the phased information of the parents’ genotype, custom python scripts were developed using the pysam library (https://github.com/pysam-developers/pysam) for efficient manipulation of the alignment data. The entire bioinformatics pipeline was implemented in the Nextflow workflow management system [30], which is available on GitHub at https://github.com/Popgen48/gvdlr/tree/main. To visualize alignment patterns in the BAM files, we used JBROWSE2 [31] and IGV [32].

### Primer design and validation of candidate variants

By examining the POLLED locus in both trios, we searched for short variants that would be specific for them. We found these putative variants: (a) 1 bp deletion (*p_ref1D_*) and a SNP with transition (*p_refT->C_*) in the HF trio (b) a SNP with transition (*P_G->A_*) in the Fleckvieh trio. For additional sequencing of the regions containing the mentioned three variants, we designed primers with a tool Primer3Plus [33]. We also sequenced both ends of the duplicated segment *P_FDUP_*. Details about PCR products are explained in Table 1. The PCR amplification was performed with 35 cycles of 30 seconds on 94 °C, 60 seconds on 59 °C and 60 seconds on 72 °C. PCR products of the trios were converted to barcoded nanopore sequencing libraries (SQK-NBD114.96) and sequenced on a Promethion R10.4.1 flowcell for approximately 2h. The resulting fastq files were aligned on the cattle reference genome sequence (ARS-UCD1.2) using minimap2 (v 2.28) with secondary alignments suppressed. Then, the bam files were visualized using Jbrowse2.

**Table 1:**
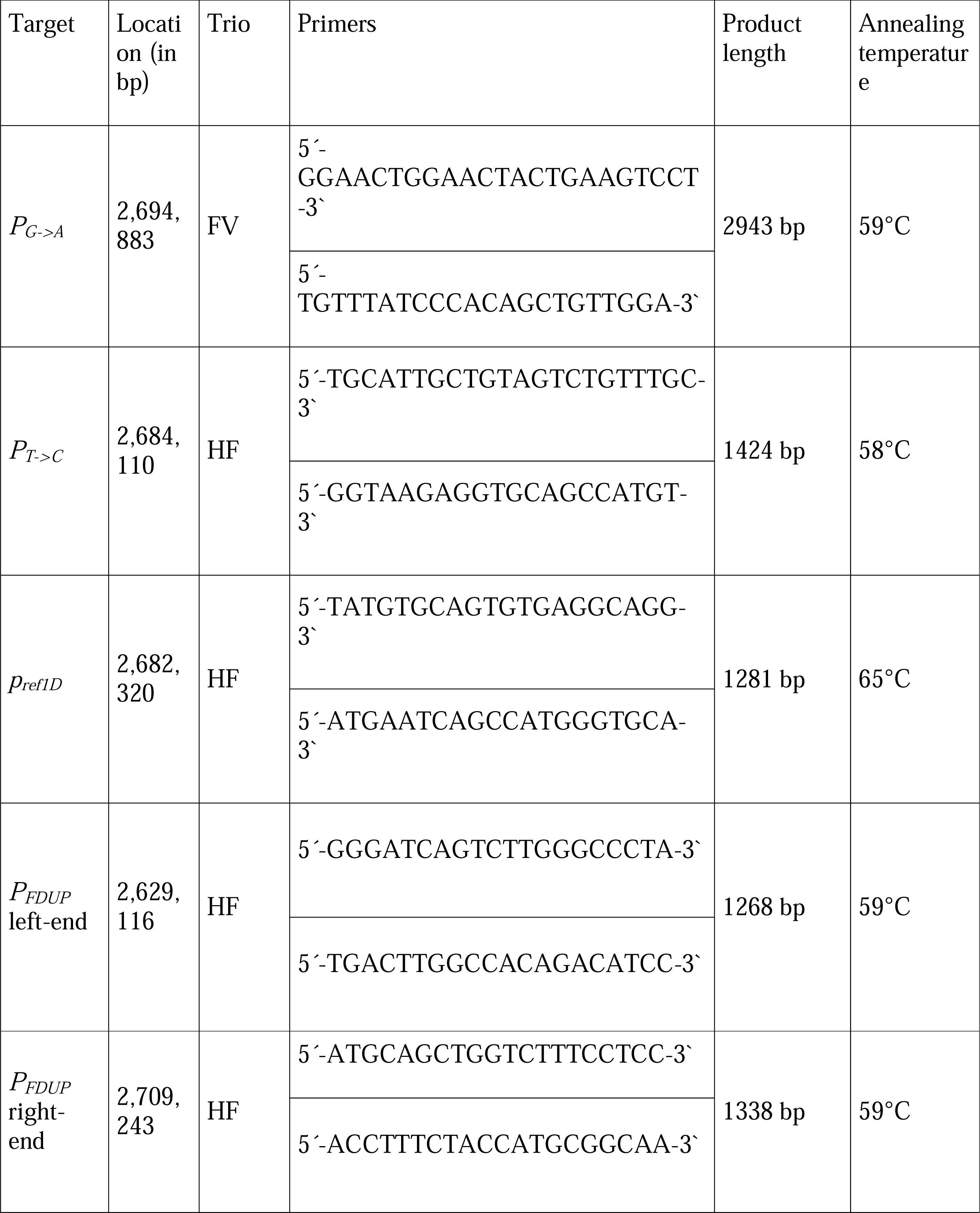
Information about PCR primers and its products used to validate candidate variants. The location refers to the position on chromosome 1 on ARS-UCD1.2 assembly.

## Results

The trios were sequenced with ONT and on average produced 36.03 Gbp sequencing data per sample with an average read length N50 of 9.70 Kbp. A detailed overview of the sequencing statistics and alignment statistics is provided in Table 2.

**Table 2.** Basic sequencing and alignment statistics of the HF and FV trio.

| Sample id | genotypes | Relationship | Total bases sequenced (in Gbp) | Total number of reads sequenced | Read length N50 (in bp) | Mean alignment coverage |
| --- | --- | --- | --- | --- | --- | --- |
| HF964 | $P_F P_F$ | Sire | 59.20 | 15,907,973 | 7,786 | 22.98x |
| HF963 | $pp$ | Dam | 25.00 | 2,670,919 | 13,986 | 9.68x |
| HF962 | $pp$ | Offspring | 59.28 | 20,506,948 | 3,957 | 22.50x |
| FV2831 | $pp$ | Sire | 18.92 | 3,564,540 | 8,379 | 7.24x |
| FV2822 | $P_F P_C$ | Dam | 31.34 | 4,180,434 | 10,001 | 12.01x |
| FV2823 | $pp$ | Offspring | 22.46 | 2,528,157 | 14,093 | 8.80x |

### Analysis of HF trio

In the HF trio, the sire was homozygous for the *P_F_* variant. This variant referred to the entire duplicated segment (Figure 1): the original segment, *p_ref_* (between 2,629,113 and 2,709,240 bp on BTA1) and its tandem duplicate (*P_FDU_*_P_). The absence of *P_F2D_* deletion in the offspring pointed towards a *de-novo* deletion of at least a part of the *P_F_* variant involving *P_FDUP_*. Concerning this deletion, five hypotheses were synthesized (Figure 2). Next, each of these five hypotheses were investigated based on the results of comprehensive variant calling and manual inspections of the entire region.

**Figure 2.**
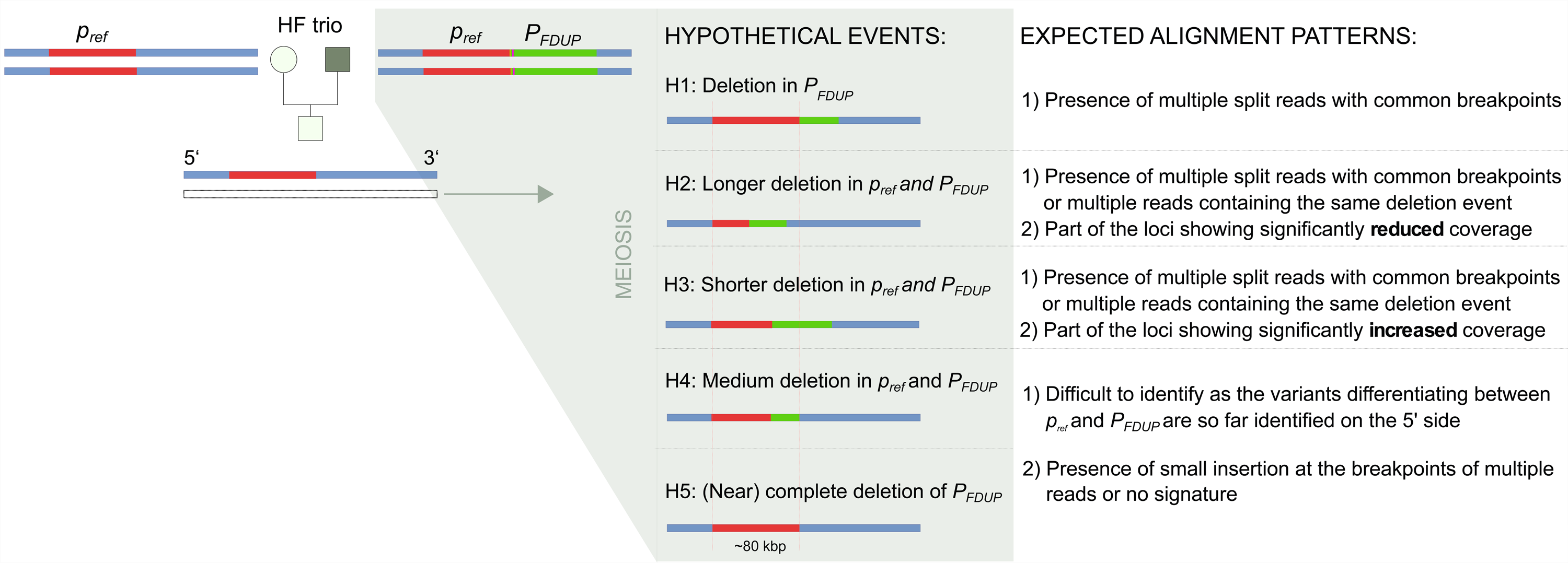
Overview of the hypotheses investigated in the HF trio. Schematic representation of the hypotheses synthesized to explain the absence of *P_F2D_* variant in the offspring of the HF trio.

### Analysis of HF sire

Using the Nextflow pipeline developed specifically for this study to identify comprehensive genomic variants from the ONT sequences, a total of two structural variants (SVs) and two SNPs were detected (Figure 1 and See Additional file 1, Table S1). Expectedly, the sire carried the *P_F_* variant (Figure 1). Interestingly, the visualization of the breakpoints identified 2 split reads that suggested that a part of the variant is possibly duplicated one more time (See Additional file 2, Figure S1). Indeed, a *de-novo* assembly of the sire also supported this observation (See Additional file 3, Figure S2).

### Analysis of HF offspring

The Nextflow pipeline identified a total of one structural variant (SV), an insertion, and eight single nucleotide polymorphisms (SNPs) in the sequences that could contain parts of segments *p_ref_* and *P_FDUP_* in the offspring. The SV (See Additional file 1, Table S1) was identified solely by the Dysgu tool; however, it was classified as a low-quality variant, as it was detected on reads with a mapping quality of 1, indicating multiple read alignments. Notably, no other tools identified any SVs supported by two or more reads. Furthermore, our custom Python tool, designed to detect split reads, did not find any regions with two or more split reads sharing common breakpoints.

Additionally, no regions larger than 1 kbp showed significantly higher or lower coverage (with more than one split read) compared to the average coverage of the region. These findings collectively suggest that the hypotheses H1, H2 and H3 (Figure 2) could be rejected.

Next to investigate the hypothesis H4 and H5, it was important to differentiate between reads that originated from *p_ref_* and those originating from *P_FDUP_*. However, apart from two variants (one SNP and a 2 bp deletion, *P_F2D_*) detected at the very beginning of the duplicated regions, no other variant that could differentiate the duplicated segment (*P_FDUP_*) from the reference sequence (*p_ref_*) has been identified in previous studies [9, 10]. To assess whether any trio-specific short variants could be used for this purpose, extensive manual inspection of the region was carried out in the bam files of sire as well as offspring. A 1 bp deletion (*p_ref1D_*, located on 2,682,320 bp) present in the alignment of both sire and offspring was identified as one such candidate (See Additional file 4, Figure S3). The long reads containing reference and alternative alleles at the candidate’s 1-bp deletion and SNP suggested their localization in the duplicated segment. However, the sequencing of PCR products to validate *p_ref1D_* and the de-novo SNP suggested that these variants were false positives (See Additional file 5, Table S2). Thus, to conclude, based on these observations, it is most likely that the genomic region showing the sequence composition of *p/p* in the offspring may have been the results of a recombination event which have happened anywhere in the *P_F_* segments and deleted almost exactly 80,128 bp of the genomic segments covering either a small part of the 3’ region of *p_ref_* and large part of the 5’ region of *P_FDUP_* segment or the entire *P_FDUP_* segment (H4 or H5 hypothesis in Figure 2). Since no insertions, duplications or multiple split reads with common breakpoints were detected in the bioinformatic and manual (window-by-window) examination of the entire 80,128 bp in the HF offspring and this offspring sequence was identical to that of an animal of the *p/p*-genotype (e.g. dam), it is obvious that the deletion in the paternal germ cell was exactly 80,128 bp long.

### Analysis of the Fleckvieh trio

In the Fleckvieh trio, the dam carried *P_C_* and *P_F_* variants each. Since the *P_F_* allele is inherited from the sire and the *P_C_* from the dam, these two variants are in *trans* phases in the homologous chromosome pair of the dam, i.e. located on different members of a homologous pair of chromosomes. Further, an allelic recombination might have resulted in recombinant haplotype (with “gene conversion”) in which the sequences roughly, until the beginning of the *P_FDUP_*, originated from the chromosome containing *P_F_* variants and the sequences after that originated from the chromosome containing *P_C_* variant (Figure 3).

**Figure 3.**
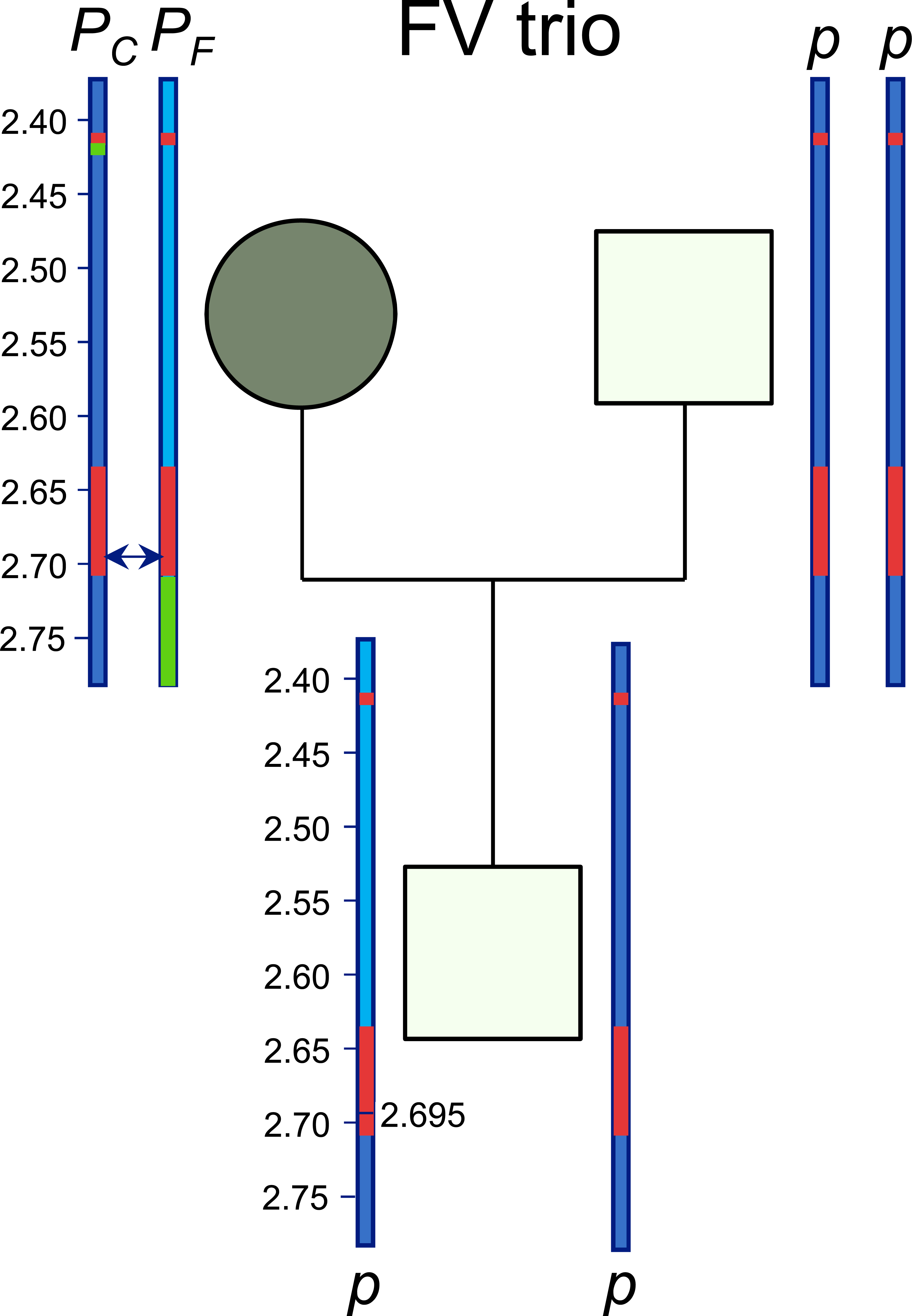
Overview of the hypothesis investigated in the Fleckvieh trio. Schematic representation of the haplotypes in the Fleckvieh trio. Note the presence of *P_F_* and *P_C_* variants in the trans-phase. The arrow between two maternal haplotypes indicates the most likely position of the recombination in the maternal germ cell.

### Analysis of Fleckvieh dam

The Nextflow pipeline identified a total of two SVs and 179 SNPs in the combined region of *P_C_* and *P_F_* variants. These two SVs were *P_C_* and *P_F_*, confirming the genotypes of the dam. Among these SVs, only *P_C_* was detected by all the tools implemented in the workflow, while *P_F_* variant was not detected by Sniffles2, one of the widely used tools to detect SV and it was detected as having low quality by cuteSV. These variants were also visually confirmed using Jbrowse2 (See Additional file 6, Figure S4 and See Additional file 7, Figure S5).

### Analysis of Fleckvieh offspring

The Nextflow pipeline identified a total of three SVs and 810 SNPs in the combined region of *P_C_* and *P_F_* variants. None of the identified SVs corresponded to the polled variants, *P_C_* and *P_F_*, confirming the results of genotyping (*pp*). The higher number of SNPs was likely due to the higher coverage (than dam); the sire also had high coverage and 910 SNPs.

The analysis of the phased SNPs in the offspring showed that one of the haplotypes, which originated from dam, had recombinant haplotype that switched from being of one parental origin (a chromosome containing *P_F_* variant) to the other (a chromosome containing *P_C_* variant) roughly around the chromosomal position, 2,694,883 bp on BTA1 (Figure 3). This resulted in skipping of both the polled variants, *P_C_* and *P_F_* in one of the haplotype of the offspring. It should be noted that although the sequencing of PCR products did not validate the SNP reported by Clair3 at this position, the switching in the haplotype was obvious from this position onwards (See Additional file 8, Table S3). This was also supported by the recombination inferred using the 50K SNP genotypes of the trio (See Additional file 9, Table S4).

## Discussion

There is growing evidence that different POLLED variants in a heterozygous state suppress horn growth to varying degrees. For example, [34] found that almost every heterozygous *P_C_/p* Fleckvieh (FV) animal forms some kind of scurs, ranging from a minimal keratin crust to a horn fused to the frontal bone. The author concluded that the visibility and strength of the scurs depends on the sex, age and polygenic background of the FV animals. Later, we compared the frequency of scurs in heterozygous *P_F_/p* and *P_C_/p* animals and confirmed that *P_F_* suppresses the development of horns significantly better than *P_C_* [35]. Furthermore, there is evidence that the *P_M_* variant completely suppresses the development of horns in heterozygous (*P_M_/p*) Mongolian Turano cattle and Mongolian yaks [11]. In our sample of 58 hornless Mongolian yaks and 40 hornless Mongolian Turano cattle, only 16 (30 %) yaks and 5 (12.5 %) Turano cattle were homozygous *P_M_/P_M_* and all others were heterozygous *P_M_/p*, but all were smooth-polled. After this observation, we tried to collect samples of Turano cattle and yaks with scurs. However, we found that breeders in Mongolia and China had never observed scurs in their animals and informed us that the animals were either horned or smooth polled. This supports our observation that the *P_M_* variant embedded in a large *P_F_*duplication could be most effective at suppressing the development of horns in heterozygous yak and cattle.

In our previous study [10], we used recombination within the *POLLED* locus to prove that segmental duplication *p_ref_-P_FDUP_* is the only remaining causal candidate for the polledness of the Friesian origin. The back-mutation of *P_F_* in the offspring of the here investigated HF trio offers the opportunity for additional confirmation of *P_F_* segmental duplication as causal variant. The HF trio was analyzed with the aim to decipher the genomic composition of the region containing the *P_F_*variant. Specifically, after observing the missing *P_F2D_* deletion in the offspring of a HF trio investigated here, we hypothesized that the most parsimonious explanation of this could be that a portion of *P_F_* in the paternal germ cell that led to the offspring must have been deleted. The deleted portion include *P_F2D_* and SNP *P_T->A_* at the transition from *p_ref_*to *P_FDUP_* (Figure 2) but we assumed that some remaining parts of *P_FDUP_* and *p_ref_* should be still present in paternal haplotype of HF offspring, i.e. one of the hypotheses H1 to H3 should apply. This would suggest that the part removed de novo in the *P_FDUP_* segment could be responsible for the polledness in cattle.

Surprisingly, our analysis neither identified any deletion nor the duplication in the entire segment in the offspring. In fact, our observations ruled out hypotheses H1, H2 and H3. The perfect reconstruction of the entire *p_ref_* segment could be achieved by a deletion of exactly 80,128 bp and then fusing of the remaining parts of *p_ref_* and *P_FDUP_* segments back to wild-type sequence (H4, Figure 2). It should also be considered that wild-type variant created in such an arrangement is indistinguishable to the structural arrangement in which the complete *P_FDUP_* sequence is deleted (H5, Figure 2): This is because *P_FDUP_* is the almost exact duplicated sequence of *p_ref_* except the two variants (*P_F2D_* and SNP *P_T->A_*) which are located at the beginning of the *P_FDUP_* segment and not present in HF offspring. Therefore, any deletion of 80,128 bp in the *P_F_* gamete, starting at any position between g.2629153 and g.2709243, will reconstruct the wild-type sequences. The only way to distinguish between H4 and H5 is to find and confirm some new variants that are present in *P_FDUP_* but lacking in the *p_ref_* segment of the HF sire or vice-versa. Our analysis did not identify any such high-quality candidate variant. Nevertheless, these results confirm that a duplication of at least part of the *p_ref_* segment (as in the *P_M_* variant) or of the entire *p_ref_* segment is required for the polled state in cattle.

Segmental duplications (SDs) are duplicated blocks of genomic DNA typically ranging in size from 1–200 kb with a sequence identity of over 90% [36]. Thus, the duplicated blocks *p_ref_* and *P_FDUP_* can be classified by definition as long and almost perfect SDs. Typically, SDs and its flanking repeats are the sites of non-allelic homologous recombination (NAHR) [37]. The direction and orientation as well as the chromosomal localization (intra-chromosomal or inter-chromosomal) of the duplicated segments involved in NAHR determine the type of SVs that may arise[37–39]. Meiotic recombination is initiated by DNA double-strand breaks (DSB). In case of allelic homologous recombination (AHR), the damaged sequence around the DSB is usually repaired by replacement with the allelic sequences of the sister chromatids as template. Mostly this recombination process is highly faithful and ensures accurate distribution of alleles and chromosomes as long as the DSB occurs in the unique regions of the genome. In the case of a DSB within the SDs regions, it is possible that the replication machinery finds the segment’s paralogs instead of the actual allelic segment for repair, resulting in NAHR [40]. All types of NAHR (intra-chromatid, inter-sister chromatid or interhomologue) can generate a deletion or other recurrent genomic rearrangements. Thus, based on the results, it is likely that, during the DSB repair in the region of *P_F_* variant, the replication machinery found the paralogous instead of allelic segment, thus resulting in hybrid SD [39]. During the process, as the repeats are very similar, it skipped the genomic region of the same size as that of the duplicated segment, resulting in a perfect deletion of one of two segments. It is noteworthy that the HF sire has more than 13,000 offspring and therefore, it could be that a gamete containing such a rare recombination event contributed to an offspring investigated here.

The polled variants *P_F_* and *P_C_* are the result of two independent mutations in different subpopulations of the bovine species. Therefore, they segregate on different haplotype backgrounds. In some breeds, such as Jersey, Fleckvieh and Holstein, both variants are inherited independently in different lines and, in rare cases, animals inherit *P_C_* from one parent and *P_F_* from another. These animals are homozygous for *POLLED*, denoted as *PP*, but heterozygous for the Celtic and for Frisian mutations, which are 200 kbp apart. The dam of the Fleckvieh trio is such an animal with *P_C_* and *P_F_* allele in trans phase. According to [41] on average 1.085 Mb corresponds to 1.0 cM in the female cattle. Therefore, on average, we expect two out of 1000 oocytes to show recombination between the *P_C_* and *P_F_* allele. It is expected that this proportion is even lower for the *POLLED* locus because in cattle all autosomal chromosomes are acrocentric [42] and regions near the centromere show the lowest recombination rate [42–44]. According to the results obtained from the analysis of the long-read sequences of FV-Trios, the following scenarios seems to be most probable: during prophase of the first meiotic division in the dam of the FV trio, a DSB was introduced in one of the sister chromatids carrying the *P_F_* variant. This most likely occurred closer to the distal end of the *p_ref_* segment ∼14 kbp upstream from the start of the *P_FDUP_* segment. This DSB was repaired by crossover with the allelic position of a non-sister chromatid containing the *P_C_* variant proximal to the DSB and *p_ref_* segment distal to the DSB. The crossover fused the proximal portion of the *P_F_*chromatid to the distal portion of the *P_C_* chromatid, thereby reconstructing the wild-type haplotype.

The results of our study point toward some interesting observations and open up intriguing possibilities that recombination events in the *POLLED* locus might result in. First, the recombination breakpoint inferred in the gametes of a dam is within 1-kb distance of *P_M_* variant. Next, in both the trios, the resulting hybrid SD has a similar composition (i.e., the wild-type arrangement was reconstructed), but are generated by different recombination types. In theory, it is equally likely that the crossover in FV-Trio fuses the *P_C_* and *P_F_* variants into a *cis*-haplotype. The offspring with *P_C_* and *P_F_* in *cis*-arrangement would be phenotypically polled and would most likely not be observed as something unexpected and therefore falsely declared as homozygous polled, *PP*. Similarly, the recombination within the SD of the *P_F_* variant could lead to diverse arrangements, which would make a reliable gene test less reliable. Therefore, it is important for large *POLLED*-locus that any deviation from the expected results is critically analyzed. Possibly some of these cases can further narrow down the sequence motive that is essential for polledness in cattle.

Our results also indicated that comprehensive detection of variants using the ONT sequences should involve application of multiple tools that implement a variety of approaches. For instance, in the FV dam, while the *P_C_* variant was identified confidently by all the tools implemented in the workflow, the *P_F_* variant was missed by Sniffles2 and detected as a low quality variant by cuteSV. Further, comprehensive detection of short variants like InDels and SNPs with high confidence are difficult using long-read sequencing technology like ONT because of its relatively high error rates. Therefore, our data suggest that the short variants identified using ONT sequencing technology should be thoroughly validated even after applying the variety of algorithms.

## Conclusions

Here, using the 50k genotyping data and long-read sequencing data generated from HF and Fleckvieh trios, we comprehensively investigated the genomic composition of Friesian and Celtic polled variants. Our data support the hypothesis that the 80-kbp duplication is the genetic variant responsible for the polled phenotype of Friesian origin. Next, we also showed that different genomic arrangements because of allelic homologous recombination and non-allelic homologous recombination in *POLLED* locus can lead to the emergence of *de-novo* ancestral horn phenotypes. These kinds of arrangements can make a reliable gene test less reliable and the derivation of the phenotype difficult to predict. Therefore, it is important for the large *POLLED*-locus that any deviation from the expected result is critically analyzed. Possibly some of these cases can further narrow down the sequence motif that is essential for polledness in cattle.

## Declarations

Not applicable

## Ethics approval and consent to participate

Not applicable

## Consent for publication

Not applicable

## Availability of data and materials

The WGS data of the trios will be deposited in the public repository upon acceptance of the manuscript. All the scripts and workflows used for the analysis can be accessed on the github page: https://github.com/Popgen48/gvdlr.

## Authors’ contributions

MU contributed to the syntheses of the hypotheses, carried out the bioinformatics analyses and wrote the first draft. AG participated in the bioinformatics analyses and edited the draft. NP designed the primers, prepared the figures and edited the draft. DS carried out the SNP genotyping, PCR validation of the variants and edited the draft. IR participated in the SNP genotyping, PCR validation of the variants and edited the draft. SK carried out WGS and edited the draft. SM provided the samples, participated in the interpretation of the results and edited the draft. IM conceived the study, proposed the hypotheses, carried out the SNP haplotyping, contributed to the first draft and figures, and coordinate the entire study. All the authors read and approved the final manuscript.

## Acknowledgements

Johann Robeis LfL Bayern and breeder of the investigated HF bull (Bokelmann GbR, Sanne, Germany) for sample taking and providing phenotypic information.

## Additional files

**Additional file 1 Table S1**

Format: word file (.docx)

Title : Table S1

Description: Summary of the comprehensive genomic variants identified using *Clair3* and *longshot* tools in the offspring of HF trio and its corresponding genotypes in its sire and dam.

**Additional file 2 Figure S1**

Format: word file (.docx)

Title : Figure S1

Description: The patterns of alignment (mainly two split reads) indicated that a part of the *P_F_* variant is duplicated multiple times in the sire of HF trio.

**Additional file 3 Figure S2**

Format: word file (.docx)

Title : Figure S2

Description: The assembled contig from the reads that were identified as split reads and supported the observation of multiple duplications in the sire of HF trio.

**Additional file 4 Figure S3**

Format: word file (.docx)

Title : Figure S3

Description: The screenshot of jbrowse showing 1 bp deletion (highlighted in pink) identified on the sire and offspring of HF trio. This deletion is inferred to be present on *p_ref_*; this inference is based on the alignment of two reads that did not have this deletion in the sire of HF trio and still covered the breakpoint without splitting. The detailed mapping information about one such read is shown.

**Additional file 5 Table S2**

Format: word file (.docx)

Title : Table S2

Description: The result of very high coverage sequencing of PCR products using ONT technology.

**Additional file 6 Figure S4**

Format: word file (.docx)

Title : Figure S4

Description: The *P_C_* variant is visually confirmed in the dam of Fleckvieh trio; the variant is shown in the three reads with the insertion block highlighted in pink color.

**Additional file 7 Figure S5**

Format: word file (.docx)

Title : Figure S5

Description: The PF variant is visually confirmed in the dam of Fleckvieh trio; the split-read alignments are seen in both the panels, referring to the tandem duplication.

**Additional file 8 Table S3**

Format: text file (.txt)

Title: Phasing and inference of recombination point in the FV trio using the variants identified from the long read sequences

Description: The tab-delimited file shows the phasing information of the variants identified in the FV dam using long-read sequences. The last two column refers to the genotypes of the offspring and sire that correspond to the respective position in the dam; the “NA” indicates that the variant was not called at this position because the sample is homozygous for the reference allele. The arrow “>” indicates the allele inherited in the offspring. Notice the switch in the position of arrows from the genomic position, 2,694,883 bp.

**Additional file 9 Table S4**

Format: word file (.docx)

Title: Phasing and inference of recombination point in the FV trio using the variants genotyped with Bovine SNP 50K array.

Description: The file shows the phased data of the FV trio using the Bovine SNP 50K array genotypes.

## References

1. Zhang Y, Huang W, Hayashi C, Gatesy J, McKittrick J. Microstructure and mechanical properties of different keratinous horns. J R Soc Interface. 2018;15:20180093.

2. Aldersey JE, Sonstegard TS, Williams JL, Bottema CDK. Understanding the effects of the bovine POLLED variants. Anim Genet. 2020;51:166–76.

3. Schafberg R, Swalve HH. The history of breeding for polled cattle. Livest Sci. 2015;179:54–70.

4. Stankowich T, Caro T. Evolution of weaponry in female bovids. Proc Biol Sci. 2009;276:4329–34.

5. Knierim U, Irrgang N, Roth BA. To be or not to be horned—Consequences in cattle. Livest Sci. 2015;179:29–37.

6. Cozzi G, Gottardo F, Brscic M, Contiero B, Irrgang N, Knierim U, et al. Dehorning of cattle in the EU Member States: A quantitative survey of the current practices. Livest Sci. 2015;179:4–11.

7. Stafford KJ, Mellor DJ. Dehorning and disbudding distress and its alleviation in calves. Vet J. 2005;169:337–49.

8. Spurlock DM, Stock ML, Coetzee JF. The impact of 3 strategies for incorporating polled genetics into a dairy cattle breeding program on the overall herd genetic merit. J Dairy Sci. 2014;97:5265–74.

9. Medugorac I, Seichter D, Graf A, Russ I, Blum H, Gopel KH, et al. Bovine polledness--an autosomal dominant trait with allelic heterogeneity. PLoS One. 2012;7:e39477.

10. Rothammer S, Capitan A, Mullaart E, Seichter D, Russ I, Medugorac I. The 80-kb DNA duplication on BTA1 is the only remaining candidate mutation for the polled phenotype of Friesian origin. Genet Sel Evol. 2014;46:44.

11. Medugorac I, Graf A, Grohs C, Rothammer S, Zagdsuren Y, Gladyr E, et al. Whole-genome analysis of introgressive hybridization and characterization of the bovine legacy of Mongolian yaks. Nat Genet. 2017;49:470–5.

12. Utsunomiya YT, Torrecilha RBP, Milanesi M, Paulan SdC, Utsunomiya ATH, Garcia JF. Hornless Nellore cattle (Bos indicus) carrying a novel 110 kbp duplication variant of the polled locus. Anim Genet. 2019;50:187–8.

13. Allais-Bonnet A, Grohs C, Medugorac I, Krebs S, Djari A, Graf A, et al. Novel Insights into the Bovine Polled Phenotype and Horn Ontogenesis in Bovidae. PLoS One. 2013;8:e63512.

14. Randhawa IAS, Burns BM, McGowan MR, Porto-Neto LR, Hayes BJ, Ferretti R, et al. Optimized Genetic Testing for Polledness in Multiple Breeds of Cattle. G3 (Bethesda). 2020;10:539–44.

15. Aldersey J, Chen T, Petrovski K, Williams J, Bottema CK. Histological characterisation of the horn bud region in 58 day old bovine fetuses. Int J Dev Biol. 2024;68:117–126.

16. Carlson DF, Lancto CA, Zang B, Kim E-S, Walton M, Oldeschulte D, et al. Production of hornless dairy cattle from genome-edited cell lines. Nat Biotechnol. 2016;34:479–81.

17. Browning BL, Tian X, Zhou Y, Browning SR. Fast two-stage phasing of large-scale sequence data. Am J Hum Genet. 2021;108:1880–90.

18. Browning BL, Zhou Y, Browning SR. A One-Penny Imputed Genome from Next-Generation Reference Panels. Am J Hum Genet. 2018;103:338–48.

19. De Coster W, D’Hert S, Schultz DT, Cruts M, Van Broeckhoven C. NanoPack: visualizing and processing long-read sequencing data. Bioinformatics. 2018;34:2666–9.

20. Li H. Minimap2: pairwise alignment for nucleotide sequences. Bioinformatics. 2018;34:3094–100.

21. Rosen BD, Bickhart DM, Schnabel RD, Koren S, Elsik CG, Tseng E, et al. De novo assembly of the cattle reference genome with single-molecule sequencing. Gigascience. 2020;9:giaa021.

22. Cleal K, Baird DM. Dysgu: efficient structural variant calling using short or long reads. Nucleic Acids Res. 2022;50:e53.

23. Jiang T, Liu S, Cao S, Wang Y. Structural variant detection from long-read sequencing data with cuteSV. Methods Mol Biol. 2022;2493:137–51.

24. Smolka M, Paulin LF, Grochowski CM, Horner DW, Mahmoud M, Behera S, et al. Publisher Correction: Detection of mosaic and population-level structural variants with Sniffles2. Nat Biotechnol. 2024;42:1616.

25. Smolka M, Paulin LF, Grochowski CM, Horner DW, Mahmoud M, Behera S, et al. Detection of mosaic and population-level structural variants with Sniffles2. Nat Biotechnol. 2024;42:1571–80.

26. Heller D, Vingron M. SVIM: structural variant identification using mapped long reads. Bioinformatics. 2019;35:2907–15.

27. Kolmogorov M, Yuan J, Lin Y, Pevzner PA. Assembly of long, error-prone reads using repeat graphs. Nat Biotechnol. 2019;37:540–6.

28. Heller D, Vingron M. SVIM-asm: structural variant detection from haploid and diploid genome assemblies. Bioinformatics. 2021;36:5519–21.

29. Lin JH, Chen LC, Yu SC, Huang YT. LongPhase: an ultra-fast chromosome-scale phasing algorithm for small and large variants. Bioinformatics. 2022;38:1816–22.

30. Di Tommaso P, Chatzou M, Floden EW, Barja PP, Palumbo E, Notredame C. Nextflow enables reproducible computational workflows. Nat Biotechnol. 2017;35:316–9.

31. Diesh C, Stevens GJ, Xie P, De Jesus Martinez T, Hershberg EA, Leung A, et al. JBrowse 2: a modular genome browser with views of synteny and structural variation. Genome Biol. 2023;24:74.

32. Robinson JT, Thorvaldsdóttir H, Winckler W, Guttman M, Lander ES, Getz G, et al. Integrative genomics viewer. Nat Biotechnol. 2011;29:24–6.

33. Untergasser A, Cutcutache I, Koressaar T, Ye J, Faircloth BC, Remm M, et al. Primer3--new capabilities and interfaces. Nucleic Acids Res. 2012;40:e115.

34. Heidrich K. Phänotypische und molekulargenetische Untersuchungen zur Vererbung von Wackelhörnern beim Fleckvieh. PhD thesis, Ludwig Maximilian University of Munich, Munich. 2018.

35. Gehrke LJ, Capitan A, Scheper C, König S, Upadhyay M, Heidrich K, et al. Are scurs in heterozygous polled (Pp) cattle a complex quantitative trait? Genet Sel Evol. 2020;52:6.

36. Vervoort L, Vermeesch JR. Low copy repeats in the genome: from neglected to respected. Explor Med. 2023;4:166–75.

37. Gu W, Zhang F, Lupski JR. Mechanisms for human genomic rearrangements. Pathogenetics. 2008;1:4.

38. Chen JM, Cooper DN, Férec C, Kehrer-Sawatzki H, Patrinos GP. Genomic rearrangements in inherited disease and cancer. Semin Cancer Biol. 2010;20:222–33.

39. Parks MM, Lawrence CE, Raphael BJ. Detecting non-allelic homologous recombination from high-throughput sequencing data. Genome Biol. 2015;16:72.

40. Sasaki M, Lange J, Keeney S. Genome destabilization by homologous recombination in the germ line. Nat Rev Mol Cell Biol. 2010;11:182–95.

41. Ma L, O’Connell JR, VanRaden PM, Shen B, Padhi A, Sun C, et al. Cattle Sex-Specific Recombination and Genetic Control from a Large Pedigree Analysis. PLoS Genet. 2015;11:e1005387.

42. Popescu PC. Chromosomes of the Cow and Bull. In: McFeely R, Popescu P, editors. Domestic Animal Cytogenetics, Adv. Vet. Sci. Comp. Med. Californina:Academic Press; 1990. p. 41–71.

43. Stapley J, Feulner PGD, Johnston SE, Santure AW, Smadja CM. Variation in recombination frequency and distribution across eukaryotes: patterns and processes. Philos Trans R Soc Lond B Biol Sci. 2017;372:20160455

44. Stapley J, Feulner PGD, Johnston SE, Santure AW, Smadja CM. Correction to ‘Variation in recombination frequency and distribution across eukaryotes: patterns and processes’. Philos Trans R Soc Lond B Biol Sci. 2018;373:20170360.

