## Additional file 1, Table S1 for "Back to the horns: a reconstruction of the ancestral horn state through distinct types of recombination events"

**Table S1**. Summary of the comprehensive genomic variants identified using *Clair3* and *longshot* tools in the offspring of HF trio and its corresponding genotypes in its sire and dam.

| Coordinates (in bp) | Ref | Alt | Genotypes | | | Tools | comment |
| --- | --- | --- | --- | --- | --- | --- | --- |
|  |  |  | offspring | sire | dam |  |  |
| 2,631,830 | A | G | 0/1 | 0/0 | 0/0 | Clair3 | Depth of alternate allele is 1 in offspring |
| 2,634,577 | T | C | 0/1 | 1/1 | 0/0 | Clair3, Longshot |  |
| 2,636,983 | T | A | 0/1 | 0/0 | 0/0 | Clair3 | Depth of alternate allele is 1 in offspring |
| 2,648,268 | G | A | 0/1 | 0/1 | 0/1 | Clair3, Longshot |  |
| 2,660,814 | G | A | 0/1 | 0/0 | 0/0 | Longshot | Most probably a de-novo SNP in the offspring |
| 2,684,110 | T | C | 0/1 | 0/0 | 0/0 | Longshot | Most probably a de-novo SNP in the offspring |
| 2,691,064 | T | C | 0/1 | 1/1 | 1/1 | Clair3, Longshot | Only one read (of the total 21 reads) in the offspring contained the reference allele |
| 2,691,336 | GC | G/- | 0/1 | 0/0 | 1/1 | Clair3 |  |
| 2,694,231 | C | AGAGTACATCATGAGAATTAAATTAAATGTGTG | 0/1 | 0/1 | 0/0 | Dysgu | The reads containing this variant has MQ1 in the offspring, indicating that the variant is most likely false positive |
