## Additional file 2, Figure S1 for "Back to the horns: a reconstruction of the ancestral horn state through distinct types of recombination events"

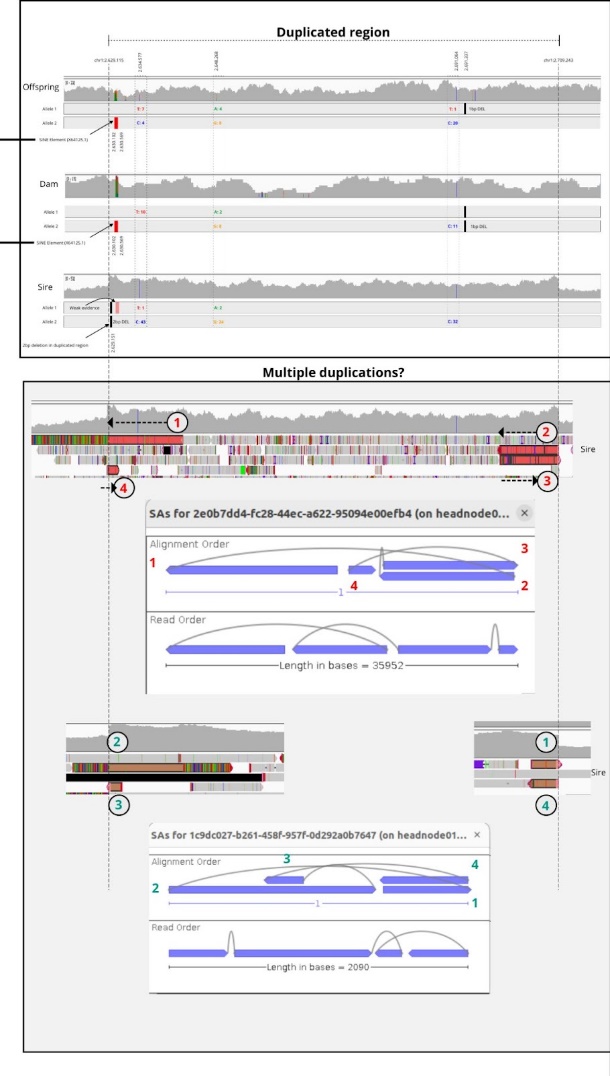

**Figure S1.** The patterns of alignment (mainly two split reads) indicated that a part of the *P_F_* variant is duplicated multiple times in the sire of HF trio.
