## Additional file 3, Figure S2 for "Back to the horns: a reconstruction of the ancestral horn state through distinct types of recombination events"

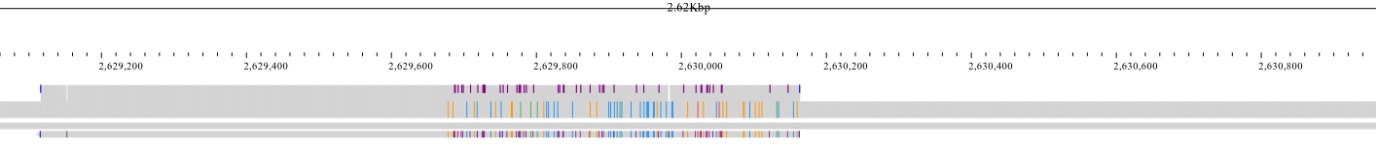

**Figure S2.** The assembled contig from the reads that were identified as split reads and supported the observation of multiple duplications in the sire of HF trio.
