## Additional file 4, Figure S3 for "Back to the horns: a reconstruction of the ancestral horn state through distinct types of recombination events"

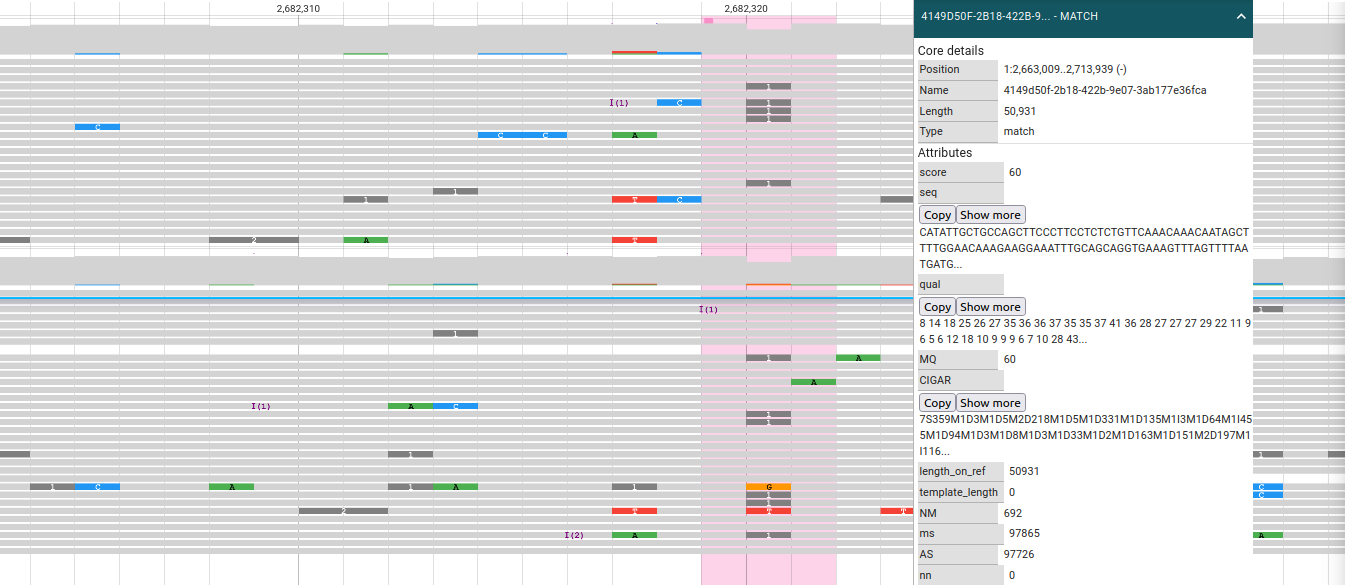

**Figure S3.** The screenshot of jbrowse showing 1 bp deletion (highlighted in pink) identified on the sire and offspring of HF trio. This deletion is inferred to be present on *p_ref_*; this inference is based on the alignment of two reads that did not have this deletion in the sire of HF trio and still covered the breakpoint without splitting. The detailed mapping information about one such read is shown.
