## Additional file 5, Table S2 for "Back to the horns: a reconstruction of the ancestral horn state through distinct types of recombination events"

**Table S2**. The result of very high coverage sequencing of PCR products using ONT technology.

| **Variant** | **Trio** | **Expected pattern** | **Observed pattern** | | | **conclusion** |
| --- | --- | --- | --- | --- | --- | --- |
|  |  |  | **sire** | **dam** | **offspring** |  |
| *P_G->A_* | FV | only dam as heterozygous | ~45,000 reads aligned, of which ~0.6% showed this SNP | ~35,000 reads aligned of which ~0.6% showed this SNP | ~11,000 reads aligned of which ~0.6% showed this SNP | False positive |
| *P_T->C_* | HF | *de-novo* in offspring | ~4098 reads aligned, of which 4% showed this SNP | ~7708 reads aligned of which 3% showed this SNP | ~11,140 reads aligned of which 4% showed this SNP | False positive |
| *p_ref1D_* | HF | Present only in sire and offspring | ~1245 reads aligned, of which 9% showed this deletion | ~2300 reads aligned, of which ~8% showed this deletion | ~4155 reads aligned, of which ~7% showed this deletion | False positive |
