## Additional file 6, Figure S4 for "Back to the horns: a reconstruction of the ancestral horn state through distinct types of recombination events"

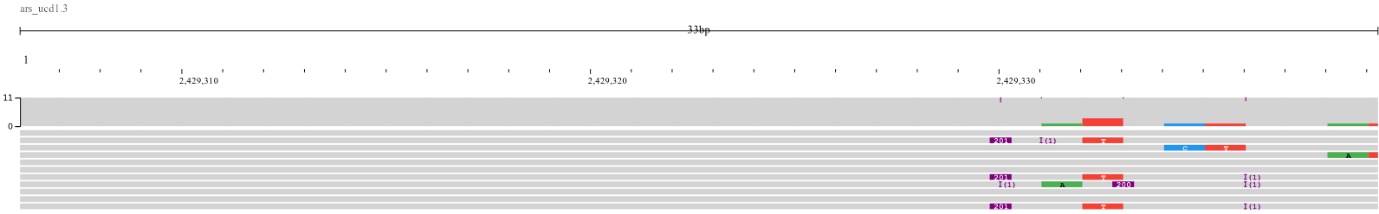

**Fig. S4.** The *P_C_* variant is visually confirmed in the dam of Fleckvieh trio; the variant is shown in the three reads with the insertion block highlighted in pink color.
