## Additional file 7, Figure S5 for "Back to the horns: a reconstruction of the ancestral horn state through distinct types of recombination events"

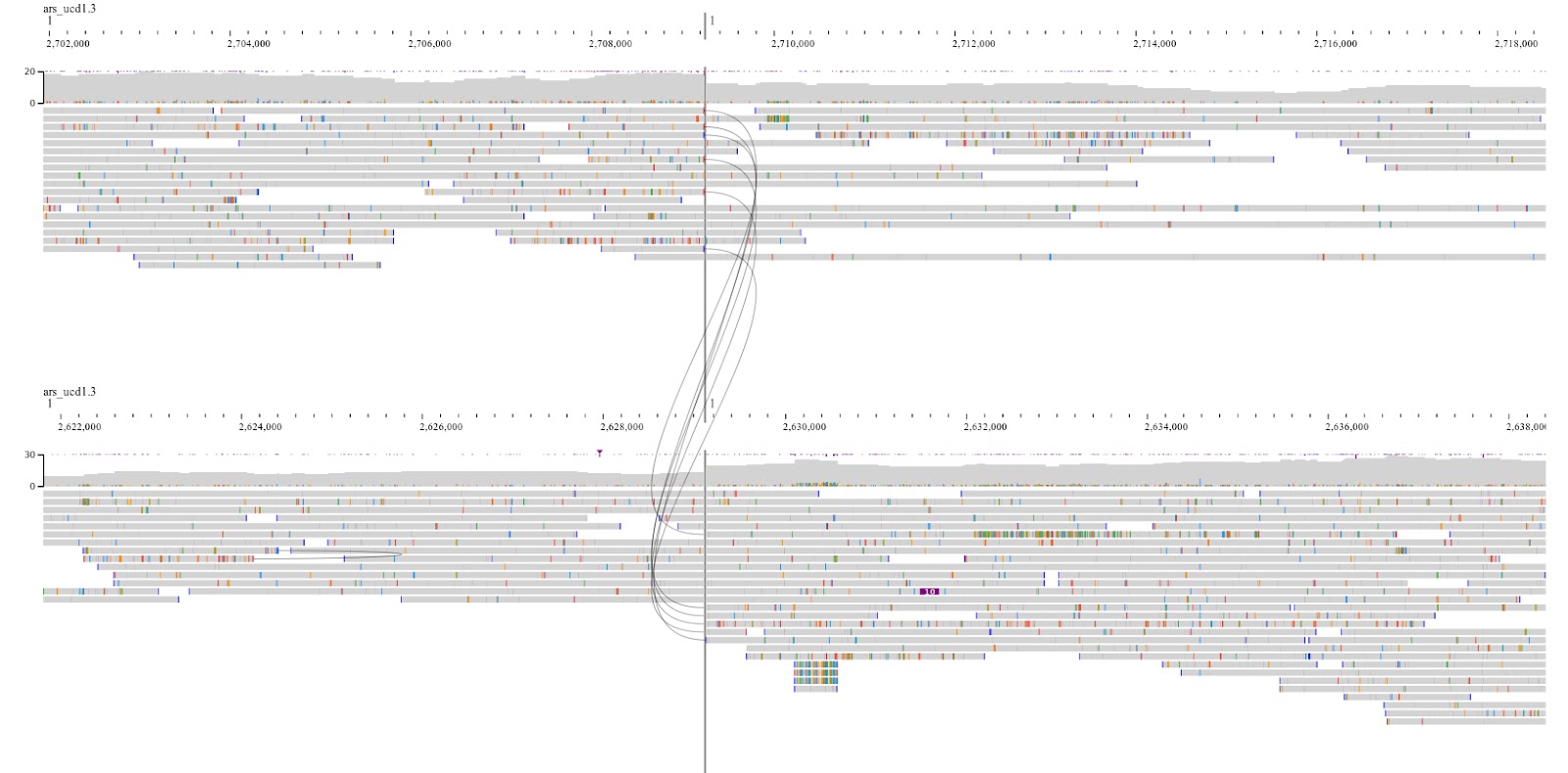

**Figure S5.** The *P_F_* variant is visually confirmed in the dam of Fleckvieh trio; the split-read alignments are seen in both the panels, referring to the tandem duplication.
