## Additional file 9, Table S4 for "Back to the horns: a reconstruction of the ancestral horn state through distinct types of recombination events"

```
AnimalLID	Nr	000000000100000000020000000003000000000400000000050000000006000000000700000000080000000009
AnimalLID	Nr	123456789012345678901234567890123456789012345678901234567890123456789012345678901234567890
FV2822K	M	GGCAGAAGGAAGGACGAGCCAGGGAGAGAGCAAAGACAAGGAAGACAGAGCAAAAGGAAAAGGAGGAGAGGGAAGGCGGGACGGAGAGC
FV2822K	P	AGCAGGGAGAAGGACGACCCGGGAAGAGGGCGAAGACAAGGAGGGCGAAGCGGCCGGGGAAGGAGGAGAAGGGCGGAGGAAAGGAGAAC
FV2823K	M	GGCAGGGGGGAGGACGACAAGAGAAAAGAGCAGCAGGGGAAGGGGCGAGGCAGCCGGGGCAGAAGAGGAAGGGCGGAGGAAAGGGGAGA
FV2823K	P	AGCAGGGAGAAGGACGACCCGGGAAGAGGGCGAAGACAAGGAAGACAGAGCAAAAGGAAAAGGAGGAGAGGGAAGGCGGGACGGAGAGC
FV2831	M	AACAGAGGGGGAGACGACCCGGGACAAGAGCAACAGGGGAAGGGGCGAGGCAGCCGGGGCAGAAGAGGAAGGGCGGAGGAAAGGGGAGA
FV2831	P	GGCAGGGGGGAGGACGACAAGAGAAAAGAGCAGCAGGGGAAGGGGCGAGGCAGCCGGGGCAGAAGAGGAAGGGCGGAGGAAAGGGGAGA
Chr	Pos	MarkerID	a1	a2
1	776231	Hapmap43437-BTA-101873	A	G
1	907810	ARS-BFGL-NGS-16466	A	G
1	1032564	Hapmap34944-BES1_Contig627_1906	A	C
1	1073496	BTA-07251-no-rs	A	G
1	1110393	ARS-BFGL-NGS-98142	A	G
1	1150763	Hapmap53946-rs29015852	A	G
1	1167048	BFGL-NGS-114208	A	G
1	1204825	ARS-BFGL-NGS-66449	A	G
1	1566539	ARS-BFGL-BAC-32770	A	G
1	1604940	ARS-BFGL-NGS-65067	A	G
1	1650662	ARS-BFGL-BAC-32722	A	G
1	1671886	ARS-BFGL-BAC-34682	A	G
1	1695632	ARS-BFGL-NGS-3964	A	G
1	1730547	ARS-BFGL-NGS-98203	A	C
1	1760809	ARS-BFGL-NGS-59105	A	C
1	1834781	ARS-BFGL-BAC-2376	A	G
1	1909740	ARS-BFGL-BAC-31722	A	G
1	1929664	BTA-49284-no-rs	C	G
1	1954528	ARS-BFGL-BAC-6557	A	C
1	1984725	ARS-BFGL-BAC-7196	A	C
1	2008865	ARS-BFGL-BAC-12931	A	G
1	2032673	ARS-BFGL-BAC-649	A	G
1	2058539	ARS-BFGL-BAC-14857	A	G
1	2082139	Hapmap53766-ss46526150	A	G
1	2107796	BFGL-NGS-115971	A	C
1	2158691	ARS-BFGL-NGS-62826	A	G
1	2235492	BTA-39394-no-rs	A	G
1	2265614	ARS-BFGL-NGS-109301	A	G
1	2305128	ARS-BFGL-NGS-105306	A	G
1	2327877	ARS-BFGL-NGS-87636	A	G
1	2348424	Hapmap24070-BTA-123581	A	C
1	2391777	ARS-BFGL-NGS-39992	A	G
1	2459049	ARS-BFGL-BAC-4401	A	G
1	2482685	ARS-BFGL-BAC-13059	A	C
1	2505294	ARS-BFGL-BAC-14930	A	G
1	2528158	ARS-BFGL-BAC-15552	A	G
1	2551347	ARS-BFGL-BAC-15580	C	G
1	2574098	ARS-BFGL-NGS-11433	A	G
1	2617150	ARS-BFGL-NGS-25183	A	G
1	2641517	ARS-BFGL-NGS-76349	A	G
1	2673226	ARS-BFGL-BAC-16253	A	G
1	2703664	ARS-BFGL-BAC-7317	A	G	=42
1	2769164	ARS-BFGL-NGS-29653	A	G	=43
1	2809407	ARS-BFGL-BAC-14982	A	G
1	2848684	BTA-120704-no-rs	A	G
1	2893133	ARS-BFGL-BAC-6733	A	C
1	2931328	ARS-BFGL-NGS-85297	A	G
1	3010910	ARS-BFGL-NGS-16727	A	G
1	3033433	BFGL-NGS-115671	A	G
1	3114153	ARS-BFGL-NGS-71085	A	G
1	3135408	ARS-BFGL-NGS-21352	A	C
1	3182687	ARS-BFGL-NGS-103391	A	G
1	3245141	ARS-BFGL-NGS-22858	A	G
1	3271366	ARS-BFGL-NGS-17308	A	C
1	3303269	ARS-BFGL-NGS-92033	A	C
1	3365623	ARS-BFGL-NGS-89044	A	G
1	3396953	ARS-BFGL-NGS-79093	A	G
1	3434539	ARS-BFGL-NGS-58892	A	G
1	3470316	ARS-BFGL-NGS-98287	A	G
1	3492221	BTB-00001612	A	C
1	3520298	Hapmap53592-rs29009975	A	G
1	3560291	ARS-BFGL-NGS-15775	A	G
1	3599891	ARS-BFGL-NGS-59192	A	G
1	3621606	ARS-BFGL-NGS-101618	A	G
1	3653664	BTB-00001299	A	G
1	3687775	ARS-BFGL-NGS-37412	A	G
1	3721639	ARS-BFGL-NGS-16784	A	G
1	3790017	BTB-02081949	A	G
1	3826765	BTB-00001074	A	G
1	3859505	ARS-BFGL-NGS-43924	A	G
1	3884274	BTA-26294-no-rs	C	G
1	3908027	ARS-BFGL-NGS-30999	A	G
1	3958736	Hapmap57114-rs29012843	A	G
1	3958867	ARS-USMARC-Parent-DQ381153-rs29012842	A	C
1	4004958	ARS-BFGL-NGS-37505	A	G
1	4052172	ARS-BFGL-NGS-107481	A	G
1	4086403	ARS-BFGL-NGS-108686	A	C
1	4113357	BTA-31687-no-rs	A	G
1	4142326	ARS-BFGL-NGS-36096	A	G
1	4168130	BTA-32603-no-rs	A	G
1	4204146	ARS-BFGL-NGS-109014	A	G
1	4250046	ARS-BFGL-NGS-69957	A	C
1	4276065	ARS-BFGL-NGS-43005	A	G
1	4309205	Hapmap59750-rs29024375	A	G
1	4335252	BTB-00002335	A	G
1	4357492	ARS-BFGL-NGS-107287	A	G
1	4394936	Hapmap42514-BTA-33574	A	G
1	4494007	ARS-BFGL-NGS-81011	A	G
1	4547357	BTB-00003162	A	C
```
